# Asexual freshwater snails make poor mate choice decisions

**DOI:** 10.1101/2021.06.08.447504

**Authors:** Sydney Stork, Joseph Jalinsky, Maurine Neiman

## Abstract

Once-useful traits that no longer contribute to fitness tend to decay over time. Here, we address whether the expression of mating-related traits that increase the fitness of sexually reproducing individuals but are likely less useful or even costly to asexual counterparts seems to exhibit decay in the latter. *Potamopyrgus antipodarum* is a New Zealand freshwater snail characterized by repeated transitions from sexual to asexual reproduction. The frequent coexistence of sexual and asexual lineages makes *P. antipodarum* an excellent model for the study of mating-related trait loss. We used a mating choice assay including sexual and asexual *P. antipodarum* females and conspecific (presumed better choice) vs. heterospecific (presumed worse choice) males to evaluate the loss of behavioral traits related to sexual reproduction. We found that sexual females engaged in mating behaviors with conspecific mating partners more frequently and for a greater duration than with heterospecific mating partners. By contrast, asexual females mated at similar frequency and duration as sexual females, but did not mate more often or for longer duration with conspecific vs. heterospecific males. These results are consistent with a scenario where selection acting to maintain mate choice in asexual *P. antipodarum* is weak or ineffective relative to sexual females and, thus, where asexual reproduction contributes to the evolutionary decay of mating-related traits in this system.

## Introduction

Adaptations that were once important for survival but subsequently become useless or impose new fitness costs tend to decay over time in a process known as vestigialization (Darwin 1859; Wiedersheim 1895; Van der Kooi and Schwander 2014). Many examples of this phenomenon can be seen throughout evolutionary history, including the loss of limbs in snakes (Lande 1978), eye structures in cave-dwelling fish (Jeffery 2005), and wings in many insect species (Roff 1990).

While the genetic mechanisms underlying vestigialization depend on evolutionary context, traits are most commonly lost by the accumulation of selectively neutral mutations (Fong et al 1995; Lahti et al 2009). The decay of presently useless characteristics exemplifies evolutionary tradeoffs: the decay of one trait frees up energy and resources to be used elsewhere by the organism (Fong et al 1995). For example, Roff (1990) suggests that the evolution of flightlessness from flighted ancestors allows for allocation of resources for reproduction. This framework implies that trait loss could translate into increased fitness for organisms for which the lost trait is no longer useful.

The decay of traits specific to mating and/or sexual reproduction (and above and beyond the loss of fitness expected under the accumulation of harmful mutations expected under asexual reproduction (Muller 1964; Hill and Robertson 1966)) is one potential consequence of transitions from sexual to asexual reproduction. These traits are typically required for reproductive success in sexual organisms. By definition, asexual organisms do not need external genetic contribution to reproduce, with the exception of sperm-dependent forms of parthenogenesis (e.g., gynogenesis). Mating behaviors are themselves associated with numerous costs (Williams 1966; Magnhagen 1991). Accordingly, vestigialization of mating-related traits in asexual lineages – including those involved in mate choice – is expected, particularly in a scenario where such behaviors and their maintenance are costly (Van der Kooi and Schwander 2014; Carson et al 1982; Kampfraath et al 2020). Alternatively, these traits may be selectively neutral or maintained by selection in asexuals, but only in a context where the costs do not outweigh the benefits of trait maintenance (Schwander et al 2013; Kraaijeveld et al 2016). Several studies have supported these ideas, finding abnormal or fully decayed traits involved in mating and reproduction (spermathecae in asexual *Timema* stick insects (Schwander et al 2013); sperm in asexual *P. antipodarum* males (Jalinsky et al 2020)) in asexual organisms. These results suggest that these traits are not being maintained by selection, at least to the extent of the same traits in sexual counterparts. Our focus is on behaviors: under the assumptions that mate preferences are associated with mate quality (Zahavi 1975; Kirkpatrick 1982; Heywood 1989), that mate quality contributes to fitness of sexual but not asexual females, that the genes underlying mating behaviors are not pleiotropic with respect to other traits also important to asexual females, and that there is at most limited gene flow between sexual and asexual conspecifics, we ask whether mating-related behaviors associated with mate preference decay in asexual females relative to sexual counterparts.

*Potamopyrgus antipodarum* is a New Zealand freshwater snail characterized by the frequent coexistence of phenotypically and ecologically similar obligately sexual and obligately asexual lineages (Lively 1987; Wallace 1992), making this species an excellent model to study the vestigialization of traits involved in sexual reproduction. Previous studies have shown that mating behavior in sexually reproducing male *P. antipodarum* does not differ from that of the presumed asexual males produced occasionally by asexual females (Soper et al 2016). While gene flow mediated by these males is formally possible (Neiman et al 2011), separate lines of evidence for distinct genomic consequences of asexuality in *P. antipodarum* (Sharbrough et al 2018; McElroy et al 2021) suggests that such gene flow is rare at most. Sexual males also show poor ability to discriminate between favorable and unfavorable mates, mating with equal frequency with sexual and asexual females and with parasitically castrated vs. healthy females (Neiman and Lively 2005). Prior emphasis on investigation of male mating behavior is largely a result of the assumption that males are physically in control of mating outcomes (Neiman and Neiman 2011), despite a lack of studies assessing the female role. These previous results set the stage for a direct comparison between sexual and asexual *P. antipodarum* females. In a mating paradigm like that demonstrated for *P. antipodarum* where males seem to lack adequate mate discrimination, differences in mating between sexual and asexual females can be attributed to mating differences between these two female groups. Therefore, we hypothesize that sexual female *P. antipodarum* are better able to choose a more favorable mate when presented with a choice than asexual counterparts.

We evaluated this hypothesis by comparing the mating behavior of sexual and asexual *P. antipodarum* females with sexual male *P. antipodarum* and sexual male *Potamopyrgus estuarinus*. *Potamopyrgus estuarinus* is a closely related and phenotypically similar species that nevertheless has minimal habitat overlap with *P. antipodarum* (Haase 2003). Sexual female *P. antipodarum* mating efforts with conspecific males are presumed to be favorable relative to mating with *P. estuarinus* because of the possibility to generate offspring only with the former. Preference for *P. antipodarum* vs. *P. estuarinus* should be selectively neutral for asexual female *P. antipodarum*, for which eggs develop in the absence of males and fertilization. We chose to use *P. estuarinus* males as a heterospecific (poor) alternative choice in this experiment because the two species are phenotypically similar and because mating behavior (but not offspring production) has been shown to occur between male and female *P. antipodarum* and *P. estuarinus* (Stork, unpublished), thereby providing *P. antipodarum* females with an adequate alternative choice.

## Materials and Methods

### Snail Selection and Care

We haphazardly selected three sets of 10 sexually mature (>3 mm long; Richards and Shinn 2004; McKenzie et al 2012; Larkin et al 2016) sexual (N=30) and asexual (N=30) female *P. antipodarum* descended from one sexual and one asexual female, respectively, collected from Lake Mapourika, New Zealand, in 2018. These six sets of 10 female snails will hereafter be referred to as ‘populations.’ We also haphazardly selected three sets of 20 each sexually mature (possessing a visible penis, Haase 2008) sexual *P. antipodarum* males (N=60) and *Potamopyrgus estuarinus* males (N=60). All *P. antipodarum* males were chosen from sexual lineages established by females sampled from New Zealand Lakes Mapourika and Selfe in 2018. These lineages were separately derived from the lineages from which we drew females. All *P. estuarinus* males were collected from the Ashley River estuary in New Zealand in February, 2020.

All snails used in the experiment had been housed in 10-L plastic tanks filled about 75% full with carbon-filtered tap water and fed *Spirulina* algae, a common *Potamopyrgus* lab food, three times weekly. These tanks were maintained in our constant-temperature snail room, which is held at 16°C and on a 12:12 hour light:dark cycle. Snails were given powdered chalk as a source of dietary calcium once weekly (feeding and maintenance following Nelson and Neiman (2011)). We isolated each population used for the experiment in a 0.95 L plastic cup containing ~700 mL of water for approximately 14 days prior to behavioral trials to control for previous exposure to males and possible recent mating. We painted female shells with light-colored nail polish to aid in differentiation between females and males when analyzing video footage. Feeding and maintenance for the populations in cups followed our standard snail room protocol.

Mating trials occurred in standard four-inch petri dishes. We covered the exterior of each dish in electrical tape to control for external visual stimuli. To begin each trial for the sexual females, we placed one sexual *P. antipodarum* female, one sexual *P. antipodarum* male, and one *P. estuarinus* male equidistant from one another around the edges of the inside of the petri dish (Fig. 1). After the three snails were placed, we filled the petri dish to about 75% total volume with carbon-filtered tap water. We used this same setup for asexual *P. antipodarum* females, with the asexual female replacing the sexual female. No snail was used in more than one trial.

**Figure 1.**
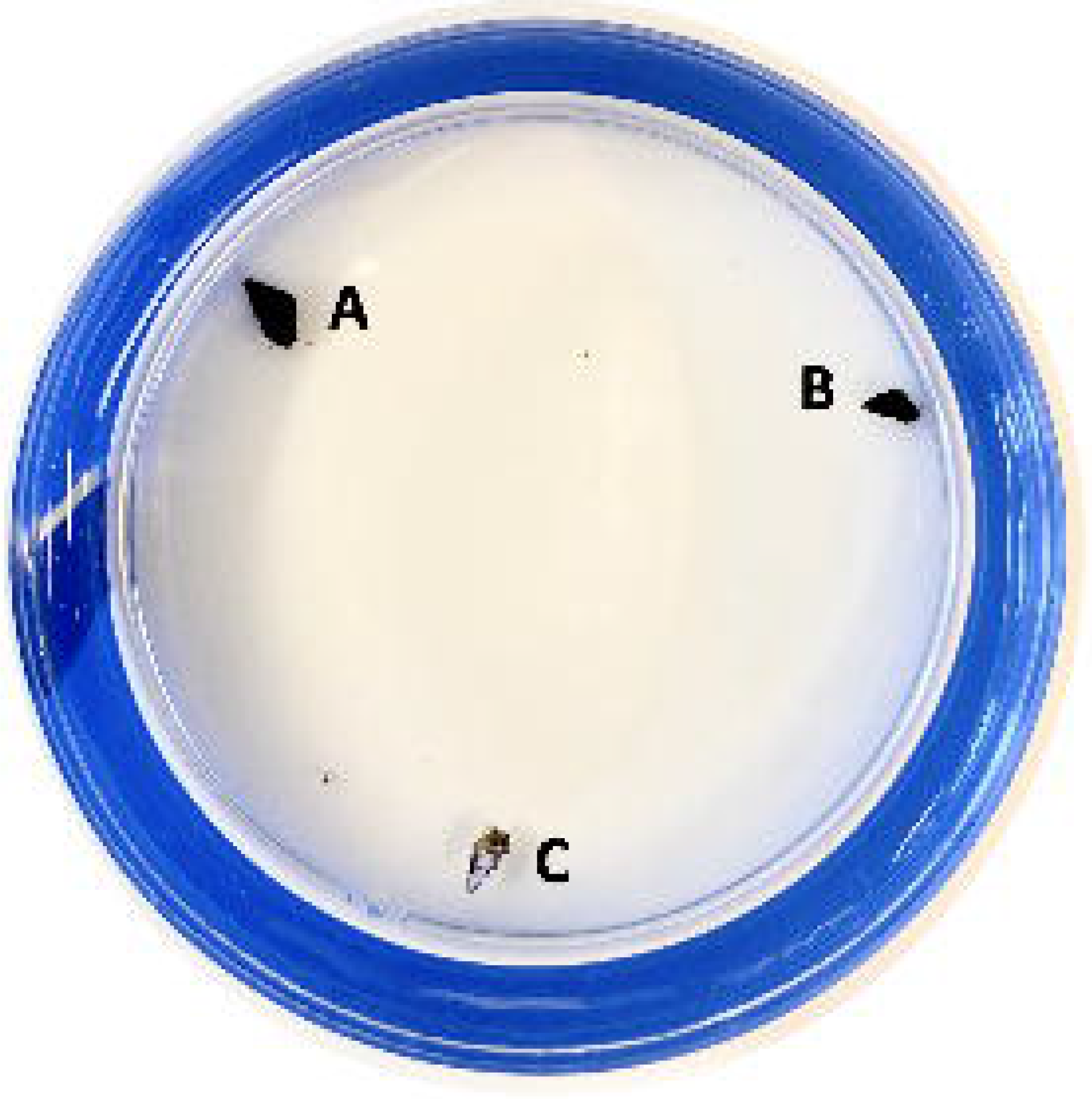
One *P. estuarinus* male (A), one *P. antipodarum* male (B), and one sexual or asexual *P. antipodarum* female (C) were placed approximately equidistant from one another around the edges of the petri dish. Each female (C) was painted a light colour with nail polish.

### Observation of Mating Behavior

We conducted each mating trial at approximately 1500 hours, for three hours. As the maximum copulation duration for *P. antipodarum* is about two hours (Neiman and Lively 2005), we established this three-hour time frame to allow for multiple mating attempts to occur. We recorded trials using a USB webcam and time-lapse video software VideoVelocity (v3.7.2090, CandyLabs 2019). The three snails per trial were allowed to move and interact freely without physical barriers inside the petri dish.

We loosely followed previous studies of mating in *P. antipodarum* (e.g., Nelson and Neiman 2011) to define attempted mating behavior as any physical interaction resulting in one snail mounting the shell of another. It is otherwise impossible to visually confirm sexual contact (joining of genitalia) in *Potamopyrgus* because this phenomenon is internal relative to the operculum and is thus obscured by the shells. Once this mounting behavior was initiated, we arbitrarily established that any such interaction lasting 15 seconds or longer would be counted as an instance of mate choice; interactions shorter than 15 seconds were potentially too short to be discernible with our time-lapse software and were excluded. Fifteen seconds is also more than enough time for snails to reorient and move on following an incidental – versus mating – encounter. Both the frequency (number of mating interactions each female had with conspecific vs. heterospecific males) and duration (length of time (seconds) that each female was in physical contact with each male type) of mating interactions between females and conspecific vs. heterospecific males were recorded.

We also used this experimental framework to establish expected outcomes and determine how these outcomes would inform our hypotheses. First, a fair test of the trait decay hypothesis would require that sexual females mate more frequently and for longer with conspecific males than with *P. estuarinus* males. This outcome is critical in indicating that sexual females seem to possess the ability to discriminate between preferential and poor mating partners in our experimental setting. Absence of a difference in mating behavior across male type for sexual females would not support the trait decay hypothesis, suggesting instead, for example, that these females are unable to differentiate between mates or that observed contact is reflective of behavior outside of mating. Mating-related trait decay in asexual females consistent with the trait decay hypothesis would be reflected in relatively weak or absent mate choice for conspecific vs. heterospecific males relative to choices in sexual female *P. antipodarum*.

### Statistical Analyses

We used the Shapiro-Wilk test to determine that both the mating frequency (sexual females: *W* = 0.769, *p* = 8.18×10^−8^; asexual females: *W* = 0.810, *p* = 1.12×10^−5^) and the mating duration (sexual females: *W* = 0.642, *p* = 1.05×10^−9^; asexual females: *W* = 0.493, *p* = 3.17×10^−8^) datasets violated normality assumptions of parametric analysis. Accordingly, we used nonparametric approaches to analyze the data. Analyses for mating duration were executed using the total mating duration for each individual (Tables S1, S2). First, we used Mann-Whitney *U* tests to compare overall mating frequency and duration between individual sexual and asexual females. Second, we used Wilcoxon rank-sum analyses to determine whether mating durations and frequencies of sexual and asexual females, respectively, with conspecific males differed from mating durations and frequencies with heterospecific males. As another means of comparing mating behavior across sexual and asexual females, we calculated discrimination scores - defined as the difference between mating frequency and mating duration, respectively, with conspecific vs. heterospecific males – for each female. We then used Mann-Whitney *U* tests to determine whether there was a significant difference in the mating frequency and mating duration discrimination scores, respectively, between sexual and asexual females. All analyses were executed with R 3.6.2 (R Core Team 2017), the ggplot2 (v3.3.3; Villanueva et al 2016), ggpubr (v0.4.0; Kassambara 2020) and dplyr (v1.0.4; Wickham et al 2021) packages.

## Results

Technical challenges with the recording software and camera hardware ultimately rendered data from three sexual female and 10 asexual female trials impossible to extract, leaving us with trials from 27 sexual females and from 20 asexual females. Comparison of the mating frequency and duration of these sexual and asexual females revealed no differences between the two female types for either trait (Frequency: Mann-Whitney *U* = 288, *z* = −0.38, *p* = 0.704; Duration: Mann-Whitney *U* = 187, *z* = 1.78, *p* = 0.075). Wilcoxon rank-sum analyses demonstrated that sexual females mated significantly more often with conspecific males (median = 1.0 matings, IQR = 2.0) than with heterospecific males (median = 1.0 matings, IQR = 1.0) (*W* = 483, *p* = 0.030) (Fig. 2a). By contrast, asexual females mated with conspecific males (median = 1.0 matings, IQR = 2.0) with the same frequency as with heterospecific males (median = 1.0, IQR = 2.5) (*W* = 172, *p* = 0.427) (Fig. 2b).

**Figure 2.**
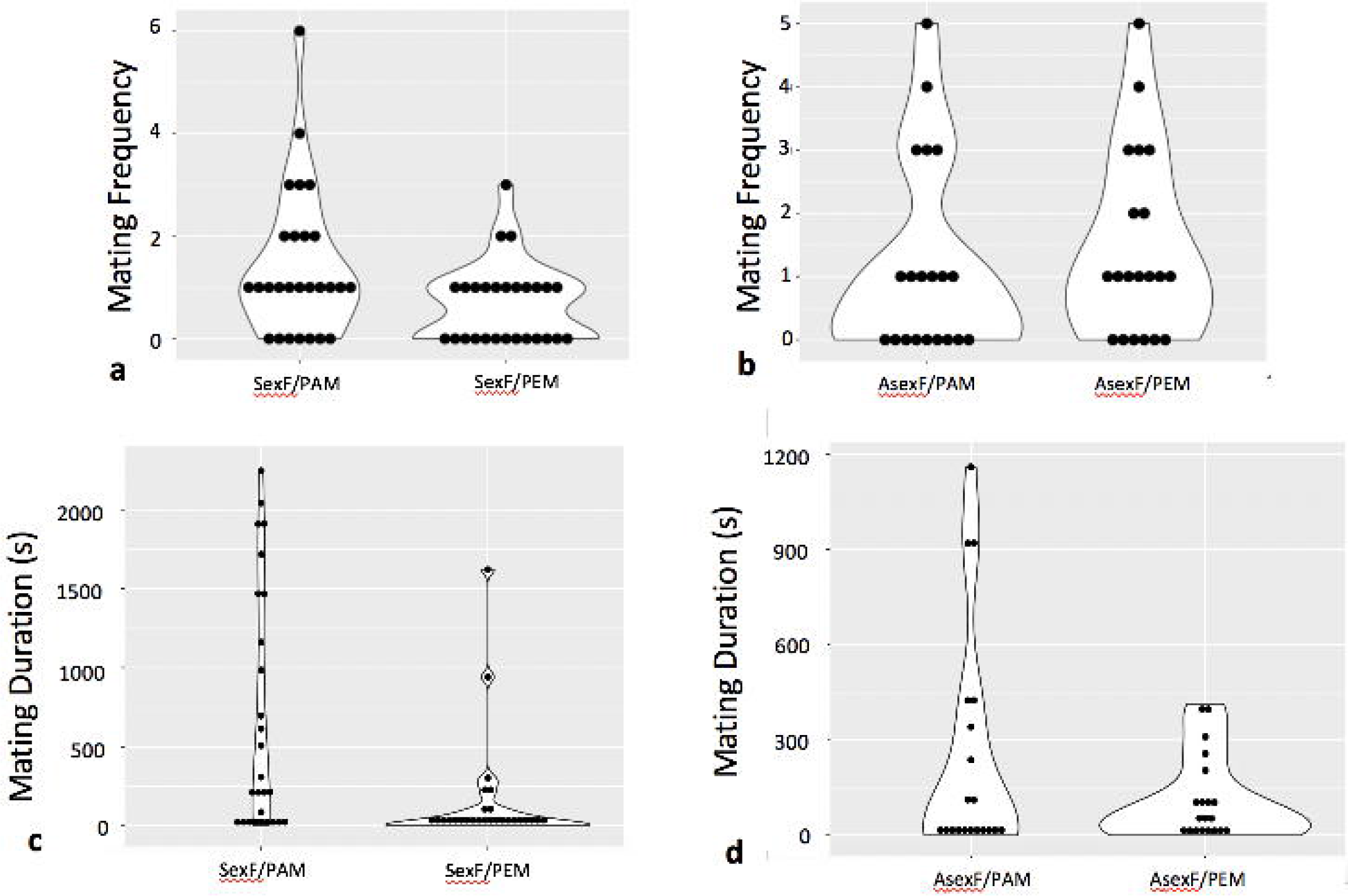
Violin plots of mating frequency (a, b) and total mating duration (c, d) of sexual (a, c; “SexF”) and asexual (b, d; “AsexF”) females with *P. antipodarum* (“PAM”) and *P. estuarinus* (“PEM”) males. Individual data points are indicated with black dots.

Sexual females also mated for significantly longer with conspecific males (median = 242.72 seconds, IQR = 1447.82) than with heterospecific males (median = 22.76 seconds, IQR = 102.25) (*W* = 520, *p* = 0.006) (Fig. 2c). Asexual females did not differ in the duration of mating with conspecific (median = 30.65 seconds, IQR = 377.43) and heterospecific males (median = 62.50 seconds, IQR = 163.84) (*W* = 202, *p* = 0.967) (Fig. 2d).

The discrimination score analysis revealed that sexual but not asexual females with a choice between conspecific and heterospecific males tend to make mating behavior decisions with respect to mating frequency that favor the former (Mann-Whitney *U* = 175.5, *z* = 2.02, *p* = 0.043). By contrast, there was no significant difference in discrimination scores for mating duration between sexual and asexual females (Mann-Whitney *U* = 208, *z* = 1.32, *p* = 0.187).

## Discussion

Our experiment revealed that sexual female *P. antipodarum* exhibited significantly higher mating frequency and longer mating duration with conspecific vs. heterospecific males. These results are as predicted if sexual female *P. antipodarum* are able to discriminate between mates of higher vs. lower quality and are important in demonstrating that discriminatory behavior can occur in the context of our experiment. By contrast, although asexual *P. antipodarum* females mated at the same frequency and for the same duration as their sexual counterparts, the asexuals showed no evidence for mate discrimination or preference. Together, these findings are consistent with a scenario where sexual but not asexual female *P. antipodarum* exhibit mate choice. Evolutionary decay of mating-related traits in an asexual context is a plausible explanation for these results and is consistent with a previous study demonstrating evidence for decay of sperm traits in the male offspring occasionally produced by asexual female *P. antipodarum* (Jalinsky et al 2020). Another non-mutually exclusive explanation for our results is provided by the possibility that asexual female *P. antipodarum* are experiencing more general degradation of phenotype associated with the accumulation of harmful mutations that is expected to accompany obligately asexual reproduction (Muller 1964; Hill and Robertson 1966). While a relatively high rate of accumulation of potentially harmful mutations has been observed in asexual *P. antipodarum* relative to sexual conspecifics (e.g., Sharbrough et al 2018), that asexual *P. antipodarum* females are similar to or even outperform sexual females when it comes to key determinants of fitness like fecundity (Paczesniak et al 2019) and growth rate and age at reproductive maturity (Larkin et al 2016) suggests that these mutations might not be translating into major phenotypic consequences. We also cannot formally exclude the possibility that asexual female snails might demonstrate mating preferences in a different context, though the fact that we observed such preferences in closely related sexual counterparts in the same experiment suggests that this explanation is not likely to entirely explain the different outcomes between sexual and asexual females. Finally, it is important to acknowledge that our study only included female *P. antipodarum* from one lake and that we only compared one sexual lineage to one asexual lineage. This Covid19-imposed limitation on our experimental design means that when possible, it will be important to repeat the experiment using additional lineages to determine whether our results are indeed more broadly generalizable.

The exhibition of similar mating behavior to their sexual counterparts combined with the lack of observable discriminatory behavior in asexual females could be explained by a situation where copulatory behavior – even in the absence of a connection between egg fertilization and embryo development – is still necessary and/or beneficial although *choice* of the mating partner is no longer important. For example, Neiman (2004) speculated that asexual females that have no direct use for sperm contributed by males might still benefit from mating if copulatory stimuli are required for maximization of reproduction, though Neiman (2006), showed that this scenario is unlikely for *P. antipodarum*. Alternatively, male ejaculate may provide nutritive benefits to asexual females. This possibility is supported by the fact that sperm storage structures are maintained in asexual *P. antipodarum* females (Dillon 2000), though evidence that these structures might also function for the digestion of waste in asexual *P. antipodarum* suggests the potential for a non-mating-specific or even unique function for this organ in asexuals (Fretter and Graham 1962). Follow-up studies of whether and how sperm/ejaculate are used in asexual *P. antipodarum* and other asexual taxa will help clarify the prevalence and persistence of copulatory behavior in asexual females.

Another and perhaps simpler explanation is that mating behavior has not fully decayed in *P. antipodarum*, and that the ability to discriminate is lost before the behavior as a whole disappears. That asexual females nevertheless engage in mating behavior at all could be a function of the relatively recent derivation of most asexual female *P. antipodarum* from sexual counterparts (Neiman et al 2005). With this in mind, it would be valuable to repeat this experiment with females from older asexual lineages (*sensu* Nelson and Neiman 2011) but departing from this prior study so that females vs. males have mate choice capabilities.

## Acknowledgments

We thank the Iowa Center for Research by Undergraduates for contributing to mentorship and training for SS, Dr. Lori Adams for her input and guidance through the Honors Program for SS; John Logsdon, Mike Winterbourn, and Mary Morgan-Richards for help in snail collection; and Marissa Roseman for setting up, characterizing, and maintaining the snail lineages that we used. Josephine Bliss also contributed to snail care. This work was supported by the Iowa Center for Research by Undergraduates, by Linda and Rick Maxson, and via NSF grant DEB-1753851. We also acknowledge very constructive and helpful critiques from Prof. Ken Kraajiveld, Prof. Ingo Schlupp, and an anonymous reviewer that greatly improved the paper.

## Declarations

### Consent to participate

Not applicable.

### Consent for publication

All authors gave final approval for publication and agree to be held accountable for the work performed therein.

### Availability of data and material

All data are available in the appendices. Snail videos are too large to readily post online but are available upon request from the authors.

### Code availability

Not applicable.

### Authors’ contributions

Sydney Stork, Joseph Jalinsky, and Maurine Neiman conceived of and designed the experiment. Sydney Stork executed the experiment and collected the data. Sydney Stork and Maurine Neiman analyzed the data. Sydney Stork created figures and drafted the manuscript. Sydney Stork, Joseph Jalinsky, and Maurine Neiman revised the manuscript.

