## Supplemental table for "Asexual freshwater snails make poor mate choice decisions"

**Supplementary Table 1 Sexual Female Mating Data**

| Snail ID | SexF/PAM Mating Frequency | SexF/PAM Mating Duration (sec) | SexF/PAM Total Mating Duration (sec) | SexF/PAM Average (mean) Mating Duration (sec) | SexF/PEM Mating Frequency | SexF/PEM Mating Duration (sec) | SexF/PEM Total Mating Duration (sec) | SexF/PEM Average (Mean) Mating Duration (sec) | Male-male Interaction Frequency | Male-male Interaction Total Duration (sec) |
| --- | --- | --- | --- | --- | --- | --- | --- | --- | --- | --- |
| 1S | 1 | 242.72 | 242.72 | 242.72 | 1 | 74.64 | 74.64 | 74.64 | 4 | 2107.72 |
| 2S | 1 | 2040.67 | 2040.67 | 2040.67 | 0 | 0.00 | 0.00 | 0.00 | 0 | 0 |
| 3S | 2 | 1899.89 | 2246.68 | 1123.34 | 0 | 0.00 | 0.00 | 0.00 | 1 | 2029.75 |
|  |  | 346.79 |  |  |  |  |  |  |  |  |
| 4S | 3 | 589.20 | 1898.68 | 632.89 | 0 | 0.00 | 0.00 | 0.00 | 2 | 3523.99 |
|  |  | 813.11 |  |  |  |  |  |  |  |  |
|  |  | 496.36 |  |  |  |  |  |  |  |  |
| 5S | 0 | 0.00 | 0.00 | 0.00 | 0 | 0.00 | 0.00 | 0.00 | 1 | 103.76 |
| 6S | 1 | 986.35 | 986.35 | 986.35 | 0 | 0.00 | 0.00 | 0.00 | 4 | 1310.08 |
| 7S | 0 | 0.00 | 0.00 | 0.00 | 1 | 102.25 | 102.25 | 102.25 | 4 | 262.44 |
| 8S | 1 | 88.29 | 88.29 | 88.29 | 1 | 942.36 | 942.36 | 942.36 | 2 | 570.7 |
| 9S | 0 | 0.00 | 0.00 | 0.00 | 0 | 0.00 | 0.00 | 0.00 | 0 | 0 |
| 10S | 0 | 0.00 | 0.00 | 0.00 | 0 | 0.00 | 0.00 | 0.00 | 0 | 0 |
| 11S | 3 | 919.00 | 1718.15 | 572.72 | 2 | 44.60 | 109.53 | 54.77 | 4 | 2420.22 |
|  |  | 767.30 |  |  |  | 64.93 |  |  |  |  |
|  |  | 31.86 |  |  |  |  |  |  |  |  |
| 12S | 2 | 337.99 | 506.98 | 253.49 | 1 | 1621.67 | 1621.67 | 1621.67 | 4 | 1118.64 |
|  |  | 168.99 |  |  |  |  |  |  |  |  |
| 13S | 1 | 183.25 | 183.25 | 183.25 | 1 | 258.80 | 258.80 | 258.80 | 0 | 0 |
| 14S | 1 | 36.71 | 36.71 | 36.71 | 1 | 64.02 | 64.02 | 64.02 | 0 | 0 |
| 15S | 3 | 136.83 | 213.59 | 71.20 | 0 | 0.00 | 0.00 | 0.00 | 5 | 3840.13 |
|  |  | 30.34 |  |  |  |  |  |  |  |  |
|  |  | 46.42 |  |  |  |  |  |  |  |  |
| 16S | 0 | 0.00 | 0.00 | 0.00 | 2 |  | 199.94 | 99.97 | 1 | 612.26 |
| 17S | 1 | 50.06 | 50.06 | 50.06 | 0 | 0.00 | 0.00 | 0.00 | 2 | 586.78 |
| 18S | 6 | 1214.21 | 1447.82 | 241.30 | 1 | 22.76 | 22.76 | 22.76 | 1 | 1423.86 |
|  |  | 38.53 |  |  |  |  |  |  |  |  |
|  |  | 63.71 |  |  |  |  |  |  |  |  |
|  |  | 31.55 |  |  |  |  |  |  |  |  |
|  |  | 25.79 |  |  |  |  |  |  |  |  |
|  |  | 74.03 |  |  |  |  |  |  |  |  |
| 19S | 1 | 306.86 | 306.86 | 308.86 | 0 | 0.00 | 0.00 | 0.00 | 2 | 915.96 |
| 20S | 1 | 1488.18 | 1488.18 | 1488.18 | 0 | 0.00 | 0.00 | 0.00 | 1 | 3200.57 |
| 21S | 0 | 0.00 | 0.00 | 0.00 | 1 | 32.77 | 32.77 | 32.77 | 2 | 344.97 |
| 22S | 1 | 236.65 | 236.65 | 236.65 | 1 | 30.95 | 30.95 | 30.95 | 1 | 4152.03 |
| 23S | 0 | 0.00 | 0.00 | 0.00 | 0 | 0.00 | 0.00 | 0.00 | 3 | 4378.67 |
| 24S | 2 | 934.78 | 1162.93 | 581.45 | 3 | 54.92 | 302.49 | 100.83 | 1 | 57.65 |
|  |  | 228.16 |  |  |  | 70.39 |  |  |  |  |
|  |  |  |  |  |  | 177.19 |  |  |  |  |
| 25S | 4 | 152.31 | 613.78 | 153.45 | 1 | 59.16 | 59.16 | 59.16 | 4 | 2244.86 |
|  |  | 182.34 |  |  |  |  |  |  |  |  |
|  |  | 143.20 |  |  |  |  |  |  |  |  |
|  |  | 136.23 |  |  |  |  |  |  |  |  |
| 26S | 1 | 696.30 | 696.30 | 696.30 | 1 | 42.78 | 42.78 | 42.78 | 3 | 2895.35 |
| 27S | 2 | 985.75 | 1922.95 | 961.48 | 0 | 0.00 | 0.00 | 0.00 | 1 | 2097.40 |
|  |  | 937.20 |  |  |  |  |  |  |  |  |

Sexual mating frequency and duration data. Each row corresponds to mating data for one individual female. Mating frequency is given in number of mating attempts. Mating duration is given in seconds. Duration data is given both for total duration (summed duration of each mating attempt) and for average duration (summed duration of each mating attempt, divided by the number of mating attempts for that individual). Analyses for mating duration were executed using the total mating duration for each individual.

**Supplementary Table 2 Asexual Female Mating Data**

| Snail ID | AsexF/PAM Mating Frequency | ASexF/PAM Mating Duration (sec) | AsexF/PAM Total Mating Duration (sec) | AsexF/PAM Average (Mean) Mating Duration (sec) | AsexF/PEM Mating Frequency | ASexF/PEM Mating Duration (sec) | AsexF/PEM Total Mating Duration (sec) | AsexF/PEM Average (Mean) Mating Duration (sec) | Male-male Interaction Frequency | Male-male Interaction Total Duration (sec) |
| --- | --- | --- | --- | --- | --- | --- | --- | --- | --- | --- |
| 1A | 1 | 341.33 | 341.33 | 341.33 | 0 | 0.00 | 0.00 | 0.00 | 5 | 4026.42 |
| 2A | 0 | 0.00 | 0.00 | 0.00 | 0 | 0.00 | 0.00 | 0.00 | 2 | 601.64 |
| 3A | 1 | 1159.90 | 1159.90 | 1159.90 | 2 | 1020.00 | 414.44 | 207.22 | 0 | 0 |
|  |  |  |  |  |  | 42.78 |  |  |  |  |
| 4A | 3 | 253.95 | 436.59 | 145.53 | 1 | 20.02 | 20.02 | 20.02 | 0 | 0 |
|  |  | 144.12 |  |  |  |  |  |  |  |  |
|  |  | 38.53 |  |  |  |  |  |  |  |  |
| 5A | 0 | 0.00 | 0.00 | 0.00 | 2 |  | 257.28 | 128.64 | 0 | 0 |
| 6A | 1 | 33.07 | 33.07 | 33.07 | 1 | 30.04 | 30.04 | 30.04 | 1 | 283.38 |
| 7A | 0 | 0.00 | 0.00 | 0.00 | 0 | 0.00 | 0.00 | 0.00 | 0 | 0 |
| 8A | 0 | 0.00 | 0.00 | 0.00 | 1 | 65.23 | 65.23 | 65.23 | 1 | 126.21 |
| 9A | 0 | 0.00 | 0.00 | 0.00 | 0 | 0.00 | 0.00 | 0.00 | 0 | 0 |
| 10A | 0 | 0.00 | 0.00 | 0.00 | 1 | 85.26 | 85.26 | 85.26 | 0 | 0 |
| 11A | 3 | 165.35 | 237.87 | 79.29 | 1 | 59.77 | 59.77 | 59.77 | 0 | 0 |
|  |  | 30.95 |  |  |  |  |  |  |  |  |
|  |  | 41.57 |  |  |  |  |  |  |  |  |
| 12A | 3 | 49.76 | 98.30 | 32.77 | 3 | 183.86 | 379.55 | 126.52 | 2 | 331.62 |
|  |  | 23.67 |  |  |  | 82.52 |  |  |  |  |
|  |  | 24.88 |  |  |  | 113.17 |  |  |  |  |
| 13A | 0 | 0.00 | 0.00 | 0.00 | 0 | 0.00 | 0.00 | 0.00 | 3 | 2210.57 |
| 14A | 1 | 28.22 | 28.22 | 28.22 | 1 | 42.78 | 42.78 | 42.78 | 2 | 2730.30 |
| 15A | 1 | 413.53 | 413.53 | 413.53 | 1 | 91.02 | 91.02 | 91.02 | 4 | 1347.10 |
| 16A | 5 | 705.10 | 903.53 | 180.71 | 3 | 71.91 | 123.18 | 41.06 | 1 | 1614.39 |
|  |  | 59.77 |  |  |  |  |  |  |  |  |
|  |  | 27.61 |  |  |  | 27.91 |  |  |  |  |
|  |  | 33.07 |  |  |  | 23.36 |  |  |  |  |
|  |  | 77.97 |  |  |  |  |  |  |  |  |
| 17A | 4 | 41.26 | 935.99 | 234.00 | 5 | 92.84 | 310.38 | 62.08 | 2 | 612.87 |
|  |  | 170.81 |  |  |  | 36.41 |  |  |  |  |
|  |  | 184.77 |  |  |  | 52.79 |  |  |  |  |
|  |  | 539.14 |  |  |  | 43.69 |  |  |  |  |
|  |  |  |  |  |  | 84.65 |  |  |  |  |
| 18A | 1 | 126.21 | 126.21 | 126.21 | 3 | 86.17 | 204.49 | 68.16 | 4 | 2829.21 |
|  |  |  |  |  |  | 74.33 |  |  |  |  |
|  |  |  |  |  |  | 43.98 |  |  |  |  |
| 19A | 0 | 0.00 | 0.00 | 0.00 | 4 | 31.86 | 115.30 | 28.82 | 0 | 0 |
|  |  |  |  |  |  | 15.47 |  |  |  |  |
|  |  |  |  |  |  | 37.93 |  |  |  |  |
|  |  |  |  |  |  | 30.04 |  |  |  |  |
| 20A | 0 | 0.00 | 0.00 | 0.00 | 0 | 0.00 | 0.00 | 0.00 | 2 | 155.95 |

Asexual mating frequency and duration data. Each row corresponds to mating data for one individual female. Mating frequency is given in number of mating attempts. Mating duration is given in seconds. Duration data is given both for total duration (summed duration of each mating attempt) and for average duration (summed duration of each mating attempt, divided by the number of mating attempts for that individual). Analyses for mating duration were executed using the total mating duration for each individual.
